# Characterisation of an inflammation-related epigenetic score and its association with cognitive ability

**DOI:** 10.1101/802009

**Authors:** Anna J. Stevenson, Daniel L. McCartney, Robert F. Hillary, Archie Campbell, Stewart W. Morris, Mairead L. Bermingham, Rosie M. Walker, Kathryn L. Evans, Thibaud S. Boutin, Caroline Hayward, Allan F. McRae, Barry W. McColl, Tara L. Spires-Jones, Andrew M. McIntosh, Ian J. Deary, Riccardo E. Marioni

## Abstract

Results from large cohort studies investigating the association between inflammation and cognition have been mixed, possibly due to methodological disparities. However, a key issue in research utilising inflammatory biomarkers is their typically phasic responses. C-reactive protein (CRP) is widely used to investigate the association between chronic inflammation and cognition, but its plasma concentrations can markedly deviate in response to acute infection. Recently a large-scale epigenome-wide association study identified DNA methylation correlates of CRP. DNA methylation is thought to be relatively stable in the short term, marking it as a potentially useful signature of exposure. Here, we generate an epigenetic CRP score and investigate its trajectories with age, and associations with cognitive ability, in comparison to serum CRP in two cohorts: a longitudinal study of older adults (the Lothian Birth Cohort 1936, n=889) and a large, cross-sectional cohort (Generation Scotland, n=7,028).

We identified differing trajectories of serum CRP across the cohorts, with no homogeneous trends seen with age. Conversely, the epigenetic score was consistently found to increase with age, and to do so more rapidly in males compared to females. Higher levels of serum CRP were found to associate with poorer cognition in Lothian Birth Cohort 1936, but not in Generation Scotland. However, a consistent negative association was identified between cognition and the epigenetic score in both cohorts. Furthermore, the epigenetic score accounted for a greater proportion of variance in cognitive ability.

Our results suggest that epigenetic signatures of acute inflammatory markers may provide an enhanced signature of chronic inflammation, allowing for more reliable stratification of individuals, and thus clearer inference of associations with incident health outcomes.

## 1. INTRODUCTION

Cognitive impairment in older age is associated with an increased risk of morbidity and mortality, and a lower quality of life (1-3). Given the generally ageing population, and the significant personal and public health burden of age-related cognitive decline, insight into its determinants and the factors contributing to individual differences is critical. The strongest known risk factor for cognitive decline is older age, suggesting a unique facet of the ageing process is likely implicated.

Epidemiological studies have associated ageing with a progressive shift to a chronic inflammatory state. This low-grade, typically sub-acute, elevation of peripheral pro-inflammatory mediators in the absence of overt infection is strongly associated with the susceptibility to, and progression of, many age-associated diseases, and is a key risk factor for mortality (4-6). Accumulating evidence has also implicated chronic systemic inflammation with incident dementia, but the association with pre-morbid age-related cognitive decline is less firmly defined, and has generated considerable debate (7). Evidence from large, prospective cohort and cross-sectional studies has been largely mixed, with positive, negative and null associations identified between serum inflammatory biomarker levels and cognitive ability (8-12).

The conflicting findings evident in the inflammation-cognition literature may be attributed to methodological disparity between studies, including marked differences in both age of participants and type of cognitive assessment batteries used, hindering inter-study comparability. However, a key issue in research utilising inflammatory biomarkers is their typically phasic responses (4). Creactive protein (CRP) – an acute-phase reactant of hepatic origin – is widely used in studies of inflammation. However, by definition, the plasma concentration of acute-phase proteins deviate by 25% or more in inflammatory disorders (13). In particular, CRP is subject to swift and considerable shifts in response to injury or acute infection. Serum levels can rapidly increase up to 1,000-fold from baseline, typically resolving to basal concentrations over a period of 7-12 days (14, 15). This poses a potential issue when utilising CRP to investigate the association between inflammation and cognition: the biological variability of CRP may not be stable enough to enable reliable stratification using blood samples from individuals gathered at a single time point (16).

DNA methylation is a widely studied epigenetic mechanism involving the addition of methyl groups to the DNA molecule, typically at Cytosine-phosphate-Guanine (CpG) dinucleotides. These modifications are involved in the regulation of gene expression and are influenced by both genetics and the environment (17). Though DNA methylation is dynamic, the short-term variability in adults is thought to be relatively stable, marking it as a potentially useful signature of chronic exposure (18-20). Through epigenome-wide association studies (EWAS), DNA methylation signals at individual CpG sites have been associated with various health and lifestyle factors, permitting the creation of methylation-based phenotypic predictors and signatures (21-24). Recently, a large-scale EWAS of serum CRP in adults identified potential DNA methylation correlates of chronic low-grade inflammation (25). Ligthart et al. identified differential methylation at 58 CpG sites that replicated across both a large European (n = 8,863) and African-American (n = 4,111) cohort. Using 7 CpG sites from this data, an inflammation-related epigenetic risk score was recently created and applied in a developmental framework investigating inflammation and child and adolescent mental health (26).

In the current study, we utilise this inflammation-related epigenetic score and characterise its relationship with serum CRP levels, its longitudinal dynamics and its association with cognitive ability in a longitudinal study of older adults (the Lothian Birth Cohort 1936) and a large, cross-sectional cohort (Generation Scotland: The Scottish Family Health Study).

## 2. METHODS

### 2.1 Lothian Birth Cohort 1936 (LBC1936)

The LBC1936 is a longitudinal study comprising individuals born in 1936, most of whom completed the Scottish Mental Survey 1947 – an intelligence test of 70,805 school children in Scotland -- aged around 11 years. Full details on the recruitment and assessment protocols of the study have been described previously (27, 28). Briefly, 1091 participants were recruited to the study aged around 70⍰years. To date, up to three further waves of testing in older age have been completed at intervals of around 3 years at mean ages of 73, 76 and 79. At each wave data has been collected on a wealth of health outcomes, lifestyle factors, cognition, and biological measures.

#### 2.1.1 DNA methylation preparation in LBC1936

Detailed information on the DNA methylation profiling of LBC1936 has been reported previously (29, 30). DNA methylation was measured at 485,512 CpG sites from whole-blood samples using the Illumina Human-Methylation450 BeadChip at the Edinburgh Clinical Research Facility. Quality control procedures were performed to remove low-quality samples (inadequate hybridisation, bisulfite conversion, nucleotide extension and staining signal) and probes with a low detection rate (<95% at p < 0.01) and low call rate (<450,000 probes detected at p < 0.01). Samples where predicted and reported sex did not match, and probes on the sex chromosomes were additionally excluded. Methylation data were available for 895 individuals at Wave 1 of the LBC1936 study.

### 2.1.2 Cognitive data in LBC1936

A general fluid-type cognitive ability score (*g*_*f*_) was derived for each participant from the first un-rotated principal component of a principal components analysis on six of the Wechsler Adult Intelligence Scale-III tests measured at Wave 2 (∼73 years). These tests assessed four different cognitive domains: letter-number sequencing and digit span backwards (working memory), digit-symbol coding and symbol search (processing speed), matrix reasoning (non-verbal reasoning) and block design (constructional ability) (31). This component explained 53% of the variance across the 6 tests, with individual test loadings ranging from 0.66 to 0.78. Full details on the testing protocol for the cognitive tests in LBC1936 has been reported previously (27, 32).

#### 2.1.3 C-reactive protein in LBC1936

Serum CRP was measured from venesected whole-blood samples. Levels were quantified using both a low-sensitivity assay (lsCRP; mg/L) at all four waves of data collection, and a high-sensitivity assay (hsCRP; mg/L) at waves 2 and 3 only. The low-sensitivity assay was performed using a dry-slide immuno-rate method on an OrthoFusion 5.1⍰F.S analyser (Ortho Clinical Diagnostics). This assay cannot distinguish values less than 3⍰mg/L, thus all readings of <3⍰mg/L were assigned a value of 1.5⍰mg/L. The high-sensitivity assay was performed at the University of Glasgow using an enzyme-linked immunosorbent assay (R&D Systems, Oxford, UK) (33).

#### 2.1.4 Genotyping in LBC1936

DNA samples were genotyped at the Edinburgh Clinical Research Facility using the Illumina 610 Quadv1 array. Preparation and quality control steps have been reported previously (34). Individuals were excluded on the basis of unresolved sex discrepancies, relatedness, and evidence of non-Caucasian descent. SNPs were included if they had a call rate ≥0.98, minor allele frequency ≥0.01, and Hardy-Weinberg equilibrium test with P ≥0.001.

### 2.2 Generation Scotland: The Scottish Family Health Study (GS)

GS is a family-based genetic epidemiology cohort sampled from the general population across three regions of Scotland. The recruitment protocol and cohort characteristics are described in detail elsewhere (35, 36). Initially, 7,953 individuals aged between 18 and 65 years were recruited between 2006 and 2011 from General Practitioner registries. Family members of these subjects aged between 18 and 99 years were then approached to participate. The final cohort comprised around 24,000 subjects (59% female). Data were collected on various cognitive, psychiatric and health measures and DNA samples were collected for genotyping and methylation profiling.

#### 2.2.1 DNA methylation preparation in GS

Genome-wide DNA methylation was profiled in samples derived from blood collected between 2006 and 2011 using the Illumina Human-MethylationEPIC BeadChip. The methylation arrays were run in two separate sets. The present study includes analysis on methylation data from 2,578 unrelated individuals in the first set (referred to herein as set 1) and 4,450 unrelated individuals in the second set (set 2). Quality control steps for both sets have been fully reported previously (37). Briefly, ShinyMethyl was used to plot the logmedian intensity of methylated versus un-methylated signal per array and outliers were excluded upon visual inspection (38). Samples in which 1% of CpGs had a detection p-value in excess of 0.05, probes with a bead count of <3 in more than 5 samples, probes in which 5% of samples had a detection p-value of >0.05, and those where predicted and recorded sex diverged were also removed.

#### 2.2.2 Cognitive data in GS

Similarly to LBC1936, *g*_*f*_ was obtained for each participant from the first un-rotated principal component of a principal components analysis of three tests of cognitive ability: logical memory, verbal fluency (executive function) and digit-symbol coding (processing speed). This component explained 49% of the variance across the three tests in set 1 and 50% in set 2. Individual test loadings in set 1 ranged from 0.61 to 0.78 and in set 2 from 0.64 to 0.77. Logical memory was assessed using the Wechsler Memory Scale III (39). Verbal fluency and digit symbol-coding were tested using the Wechsler Adult Intelligence Scale III (31). Additional information regarding the cognitive variables in GS has been described previously (40, 41).

#### 2.2.3 C-reactive protein in GS

CRP was quantified at the University of Glasgow using a commercial high-sensitivity assay on an automated analyser (c311, Roche Diagnostics, UK). Manufacturer’s calibration and quality control were employed. CRP data were available for 153 participants in Set 1 and 266 participants in Set 2. These samples had been selected as father/offspring pairs in a telomere length study.

#### 2.2.4 Genotyping in GS

Genotyping was carried out using the Illumina HumanOmniExpressExome-8 v1.0 DNA Analysis BeadChip at the Wellcome Trust Clinical Research Facility (42). The arrays were imaged on an Illumina HiScan platform and genotypes were called automatically using GenomeStudio Analysis software v2011.1. SNPs with a minor allele frequency ≤0.01 and Hardy-Weinberg equilibrium test with P < 1×10^−6^ were excluded.

### 2.3 Inflammation-related poly-epigenetic score

An inflammation-related poly-epigenetic score (i-PEGS) was derived for each participant as described by Barker et al. (26). Briefly, methylation beta values were extracted for the 7 CpG sites shown to have the strongest evidence of a functional association with serum CRP levels. These values were multiplied by their respective regression weights (corresponding to change in DNA methylation beta values per 1 unit increase in natural log-transformed CRP) taken from the largest EWAS of CRP to-date and summed to generate a single i-PEGS for each participant in each cohort (see **Supplementary Table 1** (25)). All of the regression weights from the EWAS were negative, resulting in a higher i-PEGS (ie. closer to zero) corresponding to a prediction of increased CRP levels. Of the 7 CpG sites included in the original measure, one was unavailable in the GS data (cg06126421). This CpG site was thus also excluded in the LBC1936 data and an i-PEGS inclusive of the remaining 6 CpG sites was utilised in analyses for both cohorts to allow for comparability of results. The correlation coefficient between the 7-CpG and 6-CpG scores in LBC1936 was 0.95.

**Table 1.**
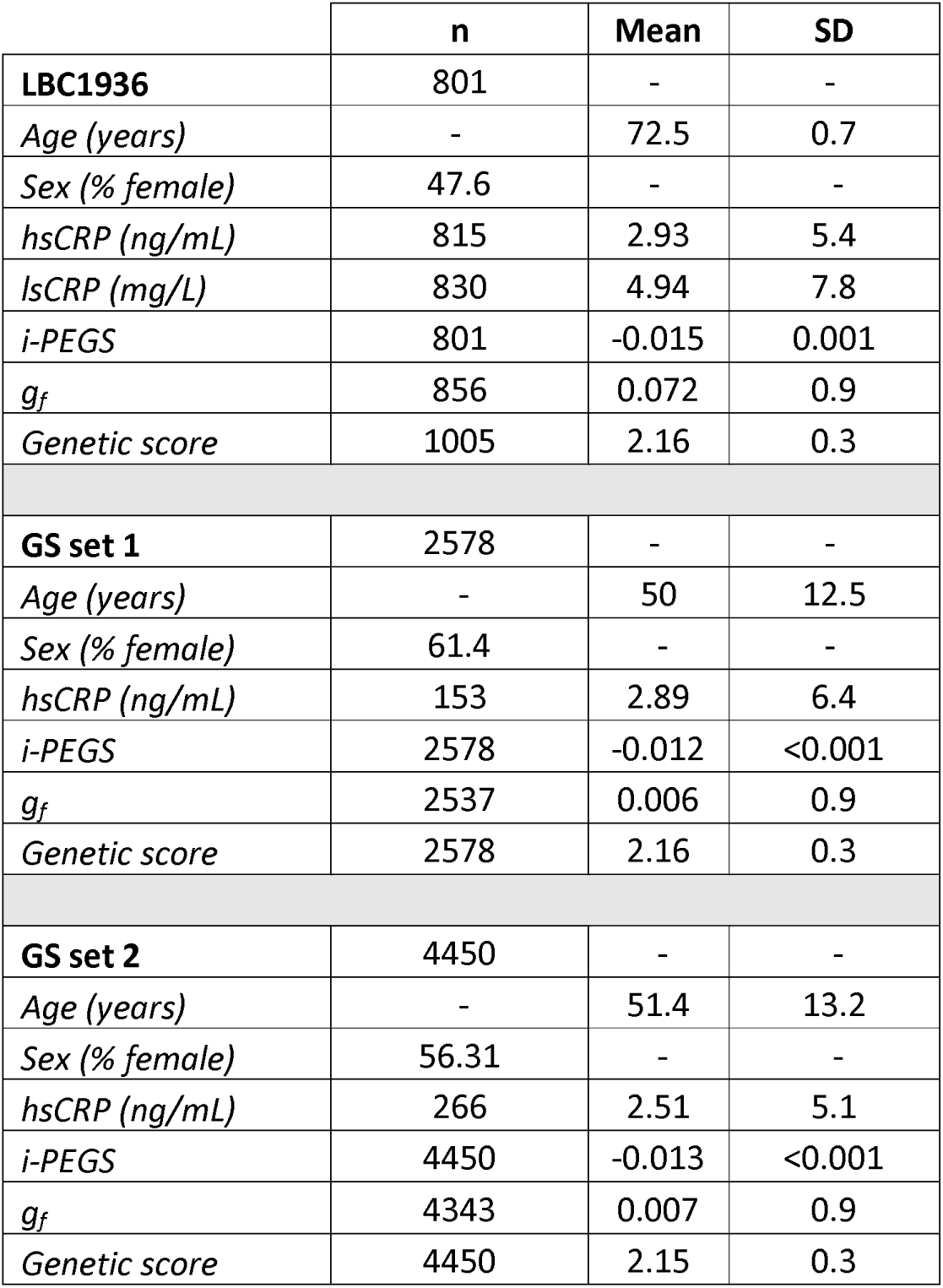
Cohort characteristics. LBC1936 summary statistics are from Wave 2 of the study. LBC1936 = Lothian Birth Cohort 1936; GS = Generation Scotland; CRP = C-reactive protein; ls = low-sensitivity, hs= high-sensitivity, i-PEGS = inflammation-related poly-epigenetic score; *g*_*f*_ = general cognitive ability score.

It should be noted that LBC1936 contributed ∼300 hsCRP samples to the EWAS of CRP from which the i-PEGS was derived (25). This may mean results from this cohort are overfitted; however the LBC1936 individuals represent a small subset of the discovery sample (n=8,863), and the probes were selected to be highly significant so it is unlikely this had a significant impact on results.

### 2.4 Genetic score for CRP

An additive weighted genetic score for CRP was constructed in both cohorts from the 18 single nucleotide polymorphisms (SNPs) that passed the genome-wide threshold (p <5×10^−8^) in the largest available GWAS of CRP to date (23). Weighted dosages were calculated by multiplying the dose of each risk allele by the effect estimate from the GWAS (see **Supplementary Table 2**). An imputation quality score of >0.8 was applied to the SNPs.

**Table 2.** Associations between *g*_*f*_ and individual predictors. LBC1936 = Lothian Birth Cohort 1936; GS = Generation Scotland; CRP = C-reactive protein; ls = low-sensitivity, hs= high-sensitivity, i-PEGS = inflammation-related poly-epigenetic score, SE = standard error.

| | Standardised $\beta$ | SE | P |
| --- | --- | --- | --- |
| **LBC1936** |  |  |  |
| *log(lsCRP)* | -0.12 | 0.03 | **0.0006** |
| *log(hsCRP)* | -0.13 | 0.04 | **0.0004** |
| *Genetic score* | 0.005 | 0.04 | 0.891 |
| *i-PEGS* | -0.14 | 0.04 | **$8.5 \times 10^{-5}$** |
| **GS set 1** |  |  |  |
| *log(CRP)* | -0.1 | 0.08 | 0.188 |
| *Genetic score* | -0.02 | 0.07 | 0.756 |
| *i-PEGS* | -0.05 | 0.02 | **0.0004** |
| **GS set 2** |  |  |  |
| *log(CRP)* | -0.02 | 0.06 | 0.698 |
| *Genetic score* | 0.05 | 0.05 | 0.313 |
| *i-PEGS* | -0.08 | 0.01 | **$1.9 \times 10^{-8}$** |

### 2.5 Statistical analysis

Pearson correlations were calculated between serum CRP, the i-PEGS and the CRP genetic score in each cohort. Inter-wave correlations and intraclass correlation coefficients were estimated for the i-PEGS and serum lsCRP over the four waves of follow-up in LBC1936 to assess the stability of the measures and their test-retest reliability.

Age- and sex-adjusted linear regression models were used to obtain the cross-sectional associations between *g*_*f*_ and serum CRP, i-PEGS and the CRP genetic score. In LBC1936 these were conducted at Wave 2 (age∼73) of the study due to the availability of both hs- and lsCRP measures at this time-point. Incremental R^2^ estimates were calculated between the null model and the models inclusive of the predictors. The R^2^ statistic represents the difference between the R^2^ of the null model (*g*_*f*_ ∼ age + sex) and the R^2^ of models with the addition of the predictors individually.

Linear mixed models were used to investigate the change in i-PEGS over the four waves available for lsCRP and 2 waves for hsCRP in LBC1936. Sex was included as a fixed effect and age (years) as the timescale. Participant ID was fitted as a random effect on the intercept. Baseline i-PEGS or serum CRP was then included as a fixed effect interaction with age to test the prediction of subsequent change in cognitive ability.

Numeric variables were scaled to have a mean of zero and a variance of 1. CRP data were log-transformed (natural log) prior to analyses due to positive skews in its distribution.

Statistical analysis was performed in R version 3.5.0 (43).

## 3. RESULTS

### 3.1 Cohort Information

Descriptive statistics of all the variables used in analyses are presented in **Table 1**. LBC1936 is an older cohort than GS (LBC1936 Wave 2: mean = 72.5 years; GS set 1: mean = 50 years; GS set 2: mean = 51.4 years), with a more even balance between sexes (LBC1936: 48% female; GS set 1: 61% female; GS set 2: 56% female). LBC1936 had the highest mean hsCRP and i-PEGS (mean hsCRP = 2.93ng/mL, mean i-PEGS = -0.015) compared to both GS set 1 (mean CRP = 2.89ng/mL, mean i-PEGS = -0.012) and set 2 (mean CRP = 2.51ng/mL, mean i-PEGS = -0.013). The mean genetic score for CRP was 2.16 in both LBC1936 and GS set 1, and 2.15 in GS set 2.

### 3.2 Correlation between serum CRP, i-PEGS and genetic score

Correlations between serum CRP, i-PEGS and the genetic score are presented in **Figure 1**. In LBC1936 and GS set 1, the correlations between the i-PEGS and serum CRP were modest (LBC1936: lsCRP: *r* = 0.25, 95% CI [0.22, 0.29]; hsCRP: *r* = 0.27, 95% CI [0.22, 0.32]; GS set 1: *r* = 0.12, 95% CI [0.04, 0.28). A stronger correlation was evident in GS set 2 (*r* = 0.39, 95% CI [0.28, 0.49]). In LBC1936 the correlation between the genetic score for CRP and serum lsCRP was 0.17 (95% CI [0.13, 0.21]). For both hsCRP in LBC1936 and GS set 2 the correlation with the genetic score was 0.26 (LBC1936: 95% CI [0.21, 0.31], GS set 2: 95% CI [0.14, 0.36]). A stronger correlation was seen in set 1 (*r* =0.35, 95% CI [0.2, 0.48]). The disparity in the correlations between cohorts may be due to there being relatively few individuals with both CRP and methylation data available in the GS cohort (set 1: n=153; set 2: n=266) and a large imbalance between males and females within these subsets (set 1: 33.9% female; set 2: 9.7% female). Additionally, there was a relatively large age difference between cohorts (LBC1936 Wave 2 mean = 72.5 years; GS set 1 mean = 46.6 years; GS set 2 mean = 55.5 years) which may have influenced results. The correlation between the i-PEGS and the genetic score was low in LBC1936 (*r* = 0.04, 95% CI [0.005, 0.08]) and slightly negative in GS (set 1: *r* = -0.03, 95% CI [-0.07, 0.008]; set 2: *r* = -0.005, 95% CI [-0.03, 0.024]).

**Figure 1.**
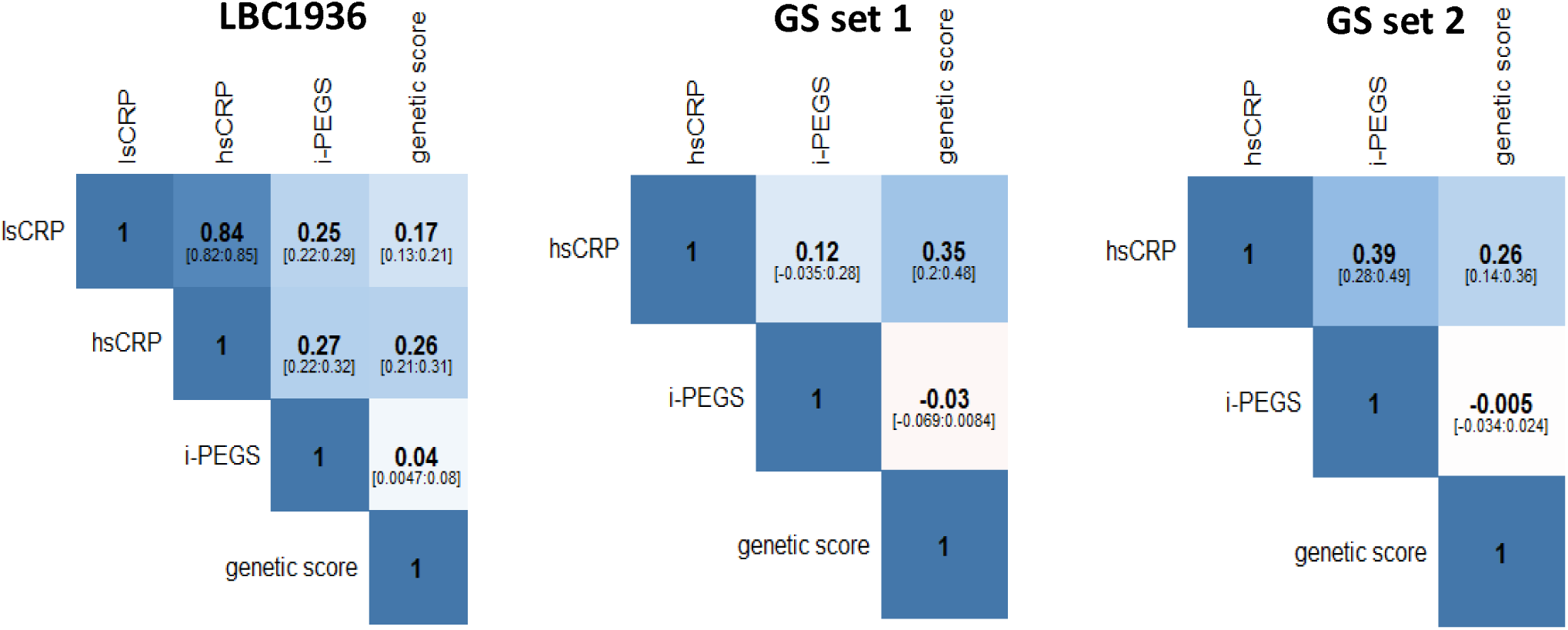
Pearson correlations between serum CRP, i-PEGS and the genetic score in LBC1936 (n=801), GS set 1 (n=153), and GS set 2 (n=266). LBC1936 = Lothian Birth Cohort 1936; GS = Generation Scotland; CRP = C-reactive protein; ls = low-sensitivity, hs= high-sensitivity, i-PEGS = inflammation-related poly-epigenetic score.

### 3.3 Trajectories, and pseudo-trajectories, of i-PEGS and serum CRP

Plots of the trajectories (pseudo-trajectories in GS) of both serum log(CRP) and i-PEGS in both cohorts are shown in **Figures 2A** and **2B**. The LBC1936 serum CRP trajectories have been plotted in a previous publication (44).

**Figure 2.**
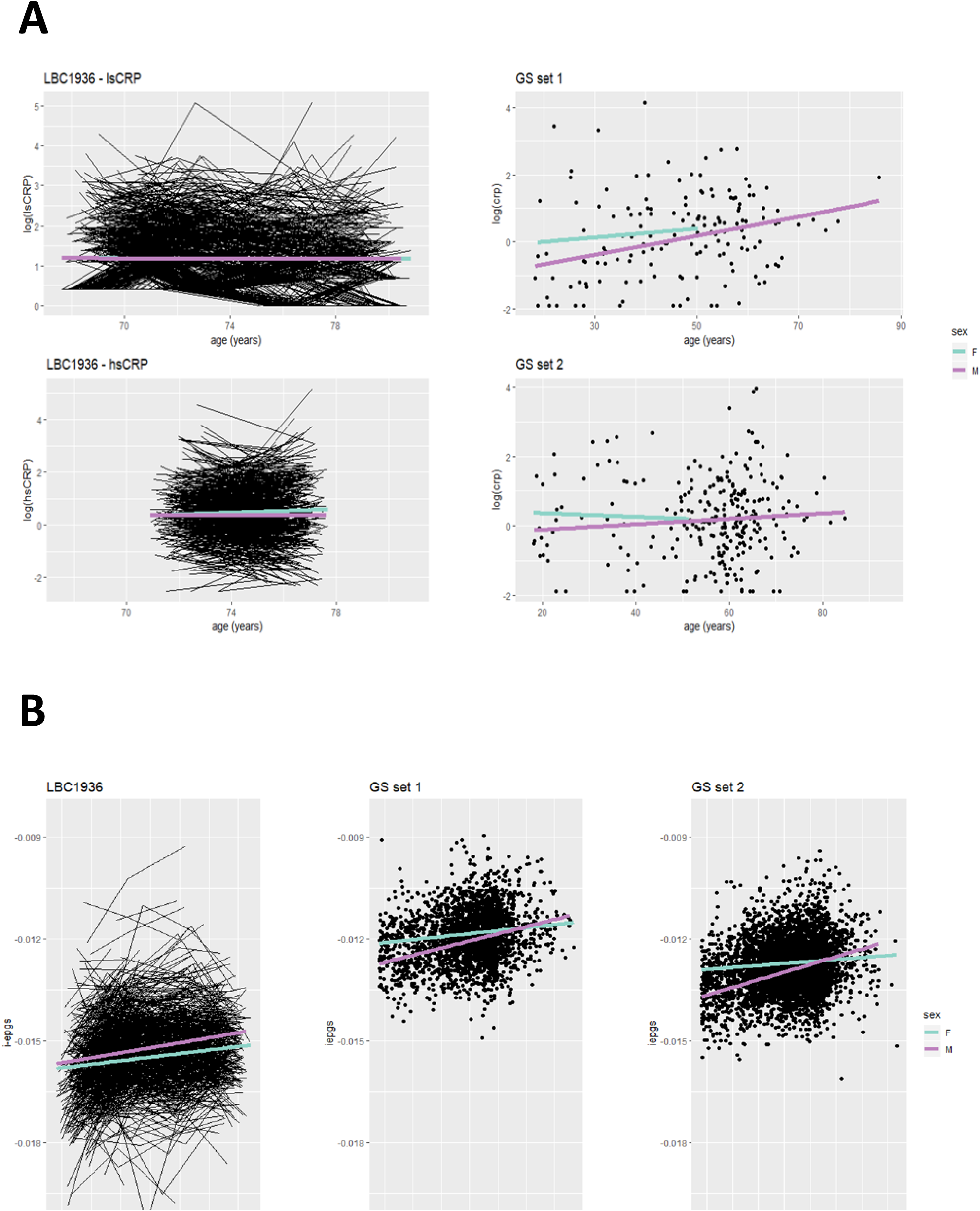
Trajectories or pseudo-trajectories of **(A)** serum CRP and **(B)** i-PEGS over age in LBC1936, GS set 1 and GS set 2. Regression lines are shown for males (purple) and females (blue). LBC1936 = Lothian Birth Cohort 1936; GS = Generation Scotland; CRP = C-reactive protein; ls = low-sensitivity, hs= high-sensitivity, i-PEGS = inflammation-related poly-epigenetic score.

The trajectories of serum CRP in LBC1936 have been reported previously (44). log(lsCRP) was found to decline over the 9 years of follow (β = -0.014, SE = 0.005, p⍰= ⍰0.004). log(hsCRP) levels measured over two waves did not significantly change (β = 0.004, SE = 0.01, p⍰= ⍰0.718). Conversely, the i-PEGS was found to increase over the 9 years of follow-up by an average of 0.07 SD per year (SE=0.004, p<2×10^−16^). A significant interaction was identified between age and sex, indicating that the i-PEGS inclined faster over time in males compared to females (β = 0.021, SE = 0.007, p = 0.004).

The intraclass correlation coefficient for lsCRP over the four waves of follow-up in LBC1936 was 0.72 (95% CI [0.69, 0.74], p<2×10^−16^). For the i-PEGS this was 0.82 (95% CI [0.75, 0.86], p<2×10^−16^), ranging from 0.60 (cg27023597) to 0.91 (cg18181703) in the 6 CpG sites that comprised the score. The correlations of the i-PEGS and serum CRP between the four waves in LBC1936 are presented in **Figure 3**. The inter-wave correlations of serum log(lsCRP) ranged from 0.35 (Wave 2-Wave 3) to 0.45 (Wave 1-Wave 2). The correlations of the i-PEGS between waves were stronger, ranging from 0.52 (Wave 1-Wave 2) to 0.71 (Wave 2-Wave 3).

**Figure 3.**
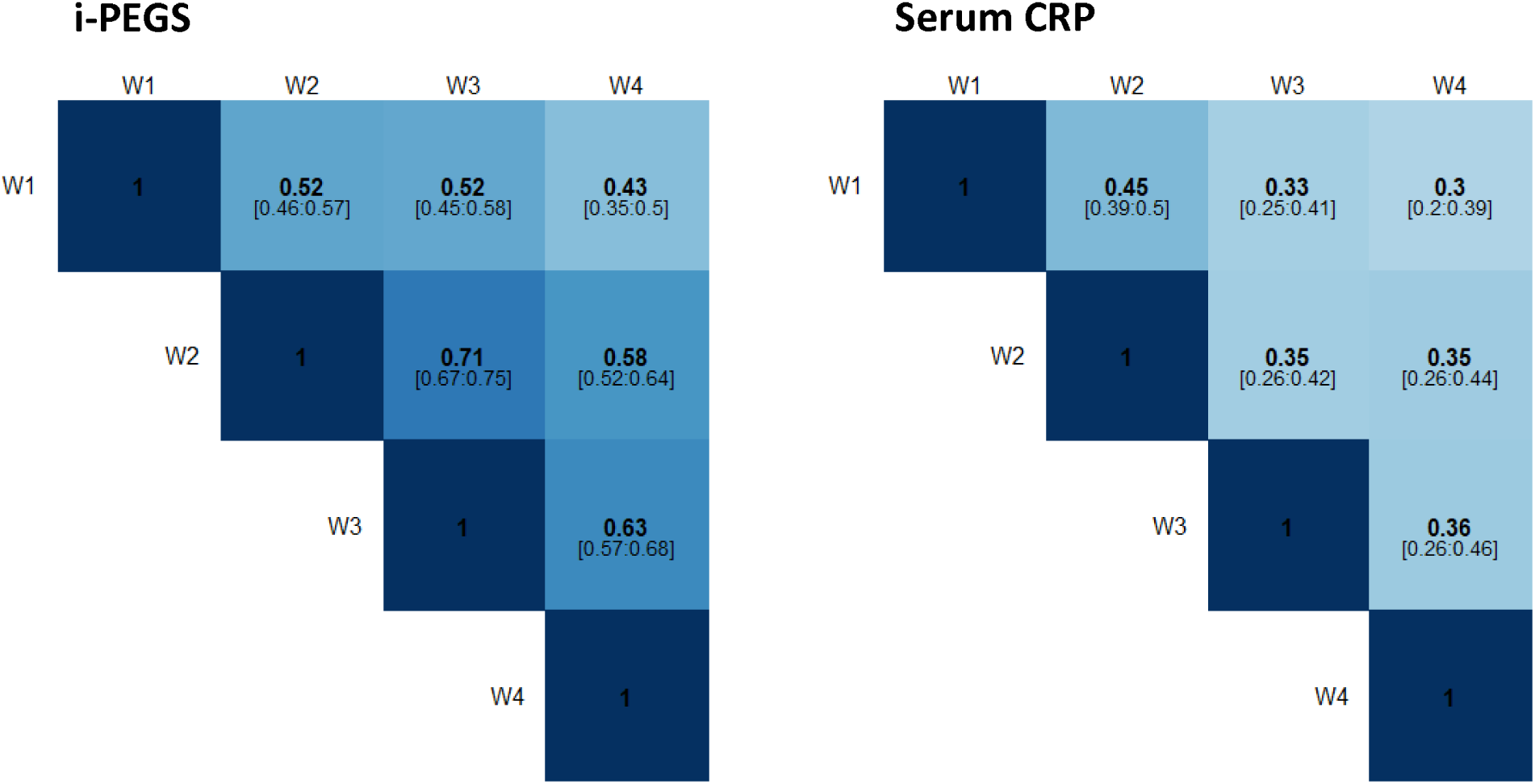
Inter-wave Pearson correlations of i-PEGS and serum CRP in LBC1936. i-PEGS = inflammation-related poly-epigenetic score; CRP = c-reactive protein; W = wave.

In GS set 1, log(CRP) significantly increased with age (β = 0.021, SE = 0.007, p = 0.002) whereas in set 2 no significant change was identified (β = 0.006, SE= 0.005, p = 0.258). In both sets, the i-PEGS increased with age with similar effect sizes (β=0.017, SE=0.001, p<2×10^−16^, in both). Again, a significant interaction between age and sex was identified, with the i-PEGS increasing more rapidly in males (set 1: β= 0.015, SE=0.003; set 2: β=0.02, SE=0.003, both p = 1.4×10^−6^)

### 3.4 Cross-sectional associations with cognitive function

The associations between the individual predictors – serum CRP, i-PEGS and the genetic score – and *g*_*f*_ are presented in **Table 2**. lsCRP has previously been found to associate with cognitive ability cross-sectionally at Wave 1 of LBC1936 (45). We found that higher CRP (both high- and low-sensitivity) was associated with lower *g*_*f*_ in LBC1936 at Wave 2 (log(lsCRP): β = -0.12, SE=0.03, p=6.0×10^−4^; log(hsCRP): β= -0.13, SE=0.04, p=3.0×10^−4^). There was no association between *g*_*f*_ and the genetic score (β= 0.005, SE=0.04, p=0.89). The i-PEGS was found to be negatively associated with *g*_*f*_ (β= -0.14, SE=0.04, p=8.5×10^−5^).

In LBC1936 the R^2^ for the null model (age and sex as the only independent variables) was 0.21. The difference between R^2^ statistics of the null model and both CRP models was similar (lsCRP: 0.013; hsCRP: 0.012). The i-PEGS explained the greatest proportion of variance with an incremental R^2^ of 0.021. An additive model inclusive of both serum CRP and the i-PEGS resulted in an incremental R^2^ 0.022.

In both GS sets, no association was identified between *g*_*f*_ and either serum CRP, or the genetic score (**Table 2**, all p≥0.188); however, higher *g*_*f*_ was significantly associated with a lower i-PEGS in both sets (set 1: β= -0.05, SE=0.02, p=4.0×10^−4^; set 2: β= -0.08, SE=0.01, p=1.9×10^−8^).

### 3.5 Longitudinal associations with cognitive function

The longitudinal associations between baseline serum CRP and i-PEGS and *g*_*f*_ in LBC1936 are presented in **Supplementary Table 3**. There was no evidence to suggest either lsCRP or i-PEGS at Wave 1 of LBC1936 was predictive of subsequent change in *g*_*f*_ over the four years of follow-up (lsCRP: p = 0.687; i-PEGS: p = 0.325).

## 4. DISCUSSION

In the present study we identified discrepant trajectories of serum CRP across the cohorts, with no homogenous trends seen with age. Conversely, a DNA methylation-based CRP score – i-PEGS – was invariably found to increase with age, and to do so more rapidly in males than in females. While we found that raised levels of serum CRP associated with poorer cognitive ability cross-sectionally, a more consistent association was identified between the latter and the i-PEGS. However, neither baseline serum CRP nor the i-PEGS were associated with longitudinal change in cognitive ability.

Research into the complex relationship between systemic inflammation and cognitive ability relies upon accurate characterisation of inflammatory mediators to enable reliable conclusions. The generally acute nature of inflammatory biomarkers - including serum CRP – mean they are particularly valuable as clinical markers for close monitoring of disease activity when repeat measures may be taken in the period of hours or days. However, in epidemiological research, analysis is typically conducted on a single blood test that may provide an imprecise picture of the true chronicity of inflammation.

While ageing is considered to be linked to systemically raised inflammation (4), we identified divergent dynamics of serum CRP in both the longitudinal (LBC1936) and cross-sectional (GS) cohorts, with increases, decreases and stable trajectories seen as a function of age. These inconsistencies highlight the challenges of utilising CRP as a biomarker of systemic inflammation in studies where it is quantified only once, or even at multiple time-points with large sampling intervals across the life-course of a longitudinal study. In contrast to serum CRP, we identified congruous increases in the i-PEGS in relation to age in both cohorts. In LBC1936, the increase per year, was greater than that seen in either GS set, potentially due to the cohort including only older individuals who may be more likely to experience a progressive elevation in inflammation (4, 46). Furthermore, we consistently found a significant interaction between i-PEGS and sex, with males having a steeper incline compared to females. This is conceivably due to men having a lower life-expectancy, and thus accelerated immune dysregulation, identified by the epigenetic score. The coherence of these results indicates the i-PEGS may be overcoming the measurement error of serum CRP, and perhaps providing a more reliable signature of chronic inflammation compared to the phenotype itself. This theory is bolstered by the higher correlations and test-retest coefficients of the i-PEGS, relative to serum CRP, between waves in LBC1936, indicating its enhanced stability over time. While the correlations between the i-PEGS and serum CRP were moderate, this may be because the single CRP measurement is not reflective of the chronic inflammatory state. As DNA methylation is considered relatively stable, the i-PEGS could conceivably be regarded as a cumulative, composite measure of inflammation akin to the HbA1c test typically utilised in diabetic patients to obtain a ∼3 month average blood glucose recording (47). The i-PEGS may then have the potential to provide a more accurate biomarker of chronic inflammation that overcomes the noise that the phasic nature of serum CRP introduces, allowing for more reliable analyses of chronic inflammatory variance and its associative relationships.

We identified significant cross-sectional associations between serum CRP and *g*_*f*_ in LBC1936. Similar associations were not seen in GS, though we may have been underpowered to detect this as relatively few participants in this cohort had CRP data available. The association between *g*_*f*_ and the i-PEGS in LBC136 was more marked than that seen with serum CRP and similar results were found in GS. Furthermore, the i-PEGS augmented the explained variance in *g*_*f*_ beyond that accounted for by serum CRP, suggesting that while the two variables are somewhat collinear, the i-PEGS harbours independent information over the raw CRP measure. Additionally, while we omitted one CpG from our analyses to allow for comparisons across cohorts, sensitivity analysis in LBC1936 using the i-PEGS inclusive of all 7 CpGs resulted in a more pronounced association with *g*_*f*_ than the 6 CpG score, suggesting these results may be conservative. No associations were found between *g*_*f*_ and the genetic score, marking the epigenetic score as having the most powerful association with cognitive ability. Our results are consistent with a recent study identifying a negative association between i-PEGS at birth, and cognitive function in early life (26). This, coupled with our results from GS, whose participant ages span early-adulthood to later-life, suggests inflammation and cognition may be related across the life-course rather than necessarily exclusively in older age. The larger effect size in LBC1936, however, suggest the association becomes more pronounced in older adults. Neither serum CRP nor the i-PEGS were associated with longitudinal change in cognitive ability over time indicating no predictive relationship between either measure of inflammation and cognitive decline.

Investigating the relationship between inflammation and cognition via the epigenetic score is particularly valuable. Firstly, it allows for research into the genomic regulation of pertinent loci, which may provide further insights into the pathways underlying the relationship, and marks them as potential therapeutic targets. Notable here is that the majority of the CpG sites included in the i-PEGS are in immune-related genes. Secondly, utilising a composite score that integrates information from multiple sites, rather than a single marker, is potentially more likely to provide a better estimate of inflammation. Finally, the epigenetic score may provide a proxy measure when CRP itself is not quantified, allowing for the investigation of inflammation in cohorts with only methylation data available.

The strengths of this study include the large sample sizes and the range of data available: GS is currently the largest epidemiological cohort studies with the availability of DNA methylation data and cognitive assessments, and LBC1936 is uniquely placed to investigate cognitive functioning, and its trajectories, in older age. The data also allowed us to investigate inflammation from a genetic, epigenetic, and phenotypic standpoint, permitting a comprehensive analysis of its relationship with cognition. However, both LBC1936 and GS are regarded as typically healthy cohorts and thus our findings may not extrapolate to the general population. It should also be noted that serum CRP is not directly produced by immune cells, and thus it is, in itself, a proxy marker of inflammation. While the i-PEGS may be capturing a more reliable picture of inflammation it would be desirable to investigate its relationship with a panel of inflammatory mediators, and to create epigenetic scores of other inflammatory mediators to test their comparative performance. Additionally, a chronic signature of inflammation might manifest in whole blood through changes in cell type proportions which should also be considered moving forward. Finally, no causal analysis was conducted in this study so it remains to be determined a) if CRP has a direct effect on methylation or indeed the opposite is true, though this has begun to be addressed elsewhere (25, 48); and b) if inflammation-related DNA methylation is causal of poorer cognition, vice-versa, or both are influenced by a confounding factor.

### Conclusion

Here, we show that an inflammation-related poly-epigenetic score may provide a more stable index of chronic, low-grade inflammation in comparison to serum CRP itself. We found the epigenetic score associated more robustly with cognitive ability and explained a greater proportion of variance compared to the measured phenotype, demonstrating the potential value in using epigenetic information in place of labile phenotypes. Epigenetic signatures of acute inflammatory markers may provide a better signature of chronic inflammation, allowing for more reliable stratification of individuals, and thus clearer inference of associations with incident health outcomes.

## ACKNOWLEDGEMENTS

The authors thank all individuals and project team members who have contributed to both GS and to the ‘STRADL: Stratifying Resilience and Depression Longitudinally’ follow-up study. GS received core support from the Chief Scientist Office of the Scottish Government Health Directorates (CZD/16/6) and the Scottish Funding Council (HR03006). Genotyping and DNA methylation profiling of the GS samples was carried out by the Genetics Core Laboratory at the Wellcome Trust Clinical Research Facility, Edinburgh, Scotland and was funded by the Medical Research Council UK and the Wellcome Trust (Wellcome Trust Strategic Award “STratifying Resilience and Depression Longitudinally” ((STRADL) Reference 104036/Z/14/Z)).

The authors thank all LBC1936 study participants and research team members who have contributed, and continue to contribute, to ongoing studies. LBC1936 is supported by Age UK (Disconnected Mind program) and the Medical Research Council (MR/M01311/1). Methylation typing was supported by Centre for Cognitive Ageing and Cognitive Epidemiology (Pilot Fund award), Age UK, The Wellcome Trust Institutional Strategic Support Fund, The University of Edinburgh, and The University of Queensland. We thank Professor Naomi Wray and Professor Peter Visscher for their useful input on the manuscript.

This work was in part conducted in the Centre for Cognitive Ageing and Cognitive Epidemiology, which is supported by the Medical Research Council and Biotechnology and Biological Sciences Research Council (MR/K026992/1) and which supports IJD. AJS and RFH are supported by funding from the Wellcome Trust 4-year PhD in Translational Neuroscience–training the next generation of basic neuroscientists to embrace clinical research [108890/Z/15/Z]. REM and DLM_C_C are supported by Alzheimer’s Research UK major project grant ARUK-PG2017B-10. TSJ and BWM are supported by the UK Dementia Research Institute which receives its funding from DRI Ltd, funded by the UK Medical Research Council, Alzheimer’s Society, and Alzheimer’s Research UK. TSJ is additionally supported by the European Research Council (ERC) under the European Union’s Horizon 2020 research and innovation programme under grant agreement No 681181.

